# Adaptive and maladaptive expression plasticity underlying herbicide resistance in an agricultural weed

**DOI:** 10.1101/2020.09.25.313650

**Authors:** Emily B. Josephs, Megan L. Van Etten, Alex Harkess, Adrian Platts, Regina S. Baucom

## Abstract

Plastic phenotypic responses to environmental change are common, yet we lack a clear understanding of the fitness consequences of these plastic responses. Here, we use the evolution of herbicide resistance in the common morning glory (*Ipomoea purpurea*) as a model for understanding the relative importance of adaptive and maladaptive gene expression responses to herbicide. Specifically, we compare leaf gene expression changes caused by herbicide to the expression changes that evolve in response to artificial selection for herbicide resistance. We identify a number of genes that show plastic and evolved responses to herbicide and find that for the majority of genes with both plastic and evolved responses, plastic responses appear to be adaptive. We also find that selection for herbicide response increases gene expression plasticity. Overall, these results show the importance of adaptive plasticity for herbicide resistance in a common weed and that expression changes in response to strong environmental change can be adaptive.

**Impact statement:** Predicting whether and how organisms will adapt to environmental change is a crucial goal. However, this goal can be complicated because environmental change can alter traits, in a process called plasticity. The extent and fitness consequences of plasticity will have important effects on the adaptive process. In this study, we use adaptation to herbicide in the agricultural weed, the common morning glory, as a model for understanding the extent and fitness consequences of plasticity in gene expression. We find evidence that gene expression plasticity is adaptive in the presence of herbicide, suggesting that understanding plasticity is crucial for understanding how organisms adapt to new environments.

## Introduction

Determining the fitness consequences of plastic responses is crucial for predicting response to selection [1], understanding the maintenance of variation for phenotypes [2], and breeding plants for new, challenging environments [3,4]. Theoretical and empirical evidence show that environments that fluctuate predictably in such a way that there is no optimum phenotype across time will favor adaptive plasticity [5,6]. Plastic responses that reduce fitness, or ‘maladaptive plasticity’, can occur when stressful environments overwhelm organisms’ ability to maintain fitness [5] or new environments expose cryptic variation, although cryptic variation can also be beneficial [7]. The presence of adaptive plasticity can allow populations to persist in the face of stressful conditions, [8] and either reduce the strength of selection by masking additive genetic variation or contribute to adaptation in novel environments [9]. Whether or not adaptive plasticity will facilitate or constrain genetic responses to selection depends on how close the plastic response gets individuals to the optimum phenotype [5]. However, despite the clear importance of disentangling adaptive from maladaptive plasticity, we lack a clear view of how the plastic responses affect fitness [5,10–12]. In particular, despite hypotheses and evidence that gene expression changes underlie many plastic responses in traditional phenotypes, the fitness consequences of gene expression plasticity are unknown [13,14].

A number of approaches have been used to describe the fitness consequences of plasticity. For example, some studies have measured fitness in organisms where a specific plastic response has or has not been induced to evaluate whether the plastic response increases fitness in the environments that induce it [15,16]. However, many plastic responses, including expression level, are not amenable to these types of manipulations. An alternative approach to understanding the fitness consequences of plasticity is comparing the fitness of genetically distinct individuals that exhibit different levels of plasticity, if there is natural genetic variation for plasticity [17–21]. In this study we focus on an alternative approach that allows us to specifically investigate gene expression variation, since it is a crucial component of plastic responses.

Here, we determine whether plastic changes in gene expression are adaptive or maladaptive by comparing plastic expression changes elicited by an environment with the expression changes that evolve during adaptation to that environment. If plasticity is adaptive and increases fitness in the new environment, the direction of plastic responses is expected to match the direction of evolved responses. Alternatively, if plasticity is maladaptive, we expect plastic responses to be opposite that of adaptive responses. Previous applications of this approach have found that plastic expression changes tend to be maladaptive in guppy brain expression response to predation environments [22] and Drosophila gene expression responses to diet [23]. An analysis of the previously mentioned guppy study along with studies from *Escherichia coli* and yeast also found a preponderance of maladaptive expression plasticity [24,25]. Strikingly, this approach has not been widely applied beyond these few studies.

We use artificially evolved glyphosate-resistant lineages of the common morning glory, *Ipomoea purpurea*, as a model for examining the fitness consequences of plastic changes in gene expression. There is a long history of using pesticide resistance evolution as models for adaptation [26,27] and the fitness costs of adaptation [27–29] but there is a significant gap in our understanding of the role of plasticity inshaping resistance evolution. Resistance to glyphosate (i.e., the active ingredient in the herbicide RoundUp) can involve either ‘target-site’ or ‘non-target site’ mutations [30]. In the former case, the genes that are the target of the herbicide contain mutations that lead to resistance whereas in the latter case any other gene that is not the target of the herbicide may be involved[27]. Non-target site resistance can be a subset of general plastic responses to abiotic stress and often involves multiple genes [31].

The common morning glory provides an excellent study system for examining questions about the role of adaptive versus maladaptive plasticity on the process of herbicide resistance evolution. Natural populations of *I. purpurea* vary in the level of glyphosate resistance [32], with non-target site herbicide resistance the most likely explanation for resistance in this species [33]. There is strong evidence of fitness costs associated with resistance in *I. purpurea*, which is consistent with the idea that resistance incurs a trade-off [34,35]. However, the specific role of plasticity in resistance evolution in *I. purpurea*, or any other weedy species is unknown.

In this study, we used seeds from an experimental evolution experiment designed to select for increased herbicide resistance in plants descending from a single population [35]. Unlike previous studies that used this approach, the population these plants were collected from did occasionally experience the new environment, herbicide treatment and displayed additive genetic variation for resistance. However, despite past experience of herbicide, artificial selection for resistance did successfully increase survival in herbicide treatments [33]. We compared leaf transcriptomes of plants from the experimentally evolved resistant lineages and from control, non-evolved lineages that were exposed to glyphosate along with replicates of each selection line that were not treated with the herbicide. We used these transcriptomes to quantify how often plastic expression responses to glyphosate aligned with the expression changes that evolved during selection for resistance. We also investigated whether selection in the herbicide environment favored increased plasticity. Overall, our results demonstrate a preponderance of adaptive gene expression plasticity in response to herbicide and that selection for increased resistance increases plasticity.

## Methods

### Study system

*Ipomoea purpurea* (L.) Roth (Convolvulaceae) is a short-lived annual vine with a relatively rapid generation time (from planted seed to producing seed within six weeks [36]) that is typically found growing on roadsides and in agricultural settings or areas of high disturbance[37]. Native to the central highlands of Mexico [38], this noxious invasive is found in every state within the US but it is particularly troublesome in the agricultural fields within the southeast and Midwestern US [32]. It has a mixed-mating system (average outcrossing rate = 0.5) with outcrossing rates that vary from highly selfing to highly outcrossing across populations [39].

### Genetic material

All plants used in the experiment descended from a single population at the previous site of the University of Georgia Plant Sciences Farm in Oconee, GA in 2000. We haphazardly sampled seeds from 122 maternal individuals at approximately every 1 meter on a transect in this population. The offspring of this sample (Generation 0 or G0) were screened for high or low glyphosate resistance in a greenhouse [40] and the offspring of the top 20% highly resistant lines (24 families) were used to be the parents for two resistant selection lines (12 parents each) and 24 parents were randomly chosen from the whole population to be the parents of two control lines. Individuals from each set of parents were grown up (Generation 1 or G1) and another generation of artificial selection was performed in the resistant selection lines using a “family selection” design by propagating siblings of the individuals in the top 20% of herbicide resistance in the population. Random individuals within each control line were chosen for the next generation to generate Generation 2 or G2 seeds. Another identical round of selection was performed on the G2 individuals to generate G3 progeny. In all crosses, individuals were used as both the pollen and ovule parent. See [35] for additional details. In a field trial, G3 plants from the resistant lines had higher survival than control lines after herbicide treatment [35].

After the field screening of G3 seeds, the 3 most resistant families were chosen from each of the selection lines and seeds from the G2 parents of these six families were crossed to each other (across lines) to generate outcrossed seeds and crossed to themselves to generate selfed seeds. The same procedure was used to generate seeds from the G2 control parents.

### Experiment

We planted 20 replicate seeds from each maternal line into two blocks and two treatments (herbicide vs no herbicide) in a randomized block design in a fenced one acre agricultural field at the Matthaei Botanical Gardens (MBGNA) at the University of Michigan in the spring of 2015. There are no natural *I. purpurea* individuals anywhere on or surrounding MBGNA. Throughout the experiment, we watered plants when needed to prevent wilting. After 6 weeks of growth, we applied 0.84 kg ai/ha glyphosate (slightly higher than ½ the suggested field dose of 1.54 kg ai/ha) to the plants in the herbicide treatment with a CO_2_ sprayer. We used a relatively low level of herbicide to cause stress but avoid killing the plants.

We collected leaf tissue from seeds at two time points post-herbicide application: 8 and 32 hours after spraying. Within each time point we randomly chose two individuals per family (one replicate individual generated from selfing and one from outcrossing per maternal line) and froze 1-2 young leaves in liquid nitrogen. All tissue at a given time point was collected and frozen within one hour.

### Transcriptome data generation

We extracted RNA from leaf tissue with Qiagen RNAeasy kits. We individually indexed libraries using the llumina TRUseq96 indexer mRNA stranded kit. Pooled libraries were run on 7 lanes of 50bp, single end sequencing on the Illumina HiSeq 4000 resulting in an average of 28 million reads per individuals.

We aligned single-end (1×52nt) adapter-trimmed Illumina RNA-seq reads separately for all samples to the Ipomoea purpurea v 2.0 genome [41] using the splice aware STAR aligner in its basic two-pass mode [42]. Genome annotation for STAR was generated using GATK’s CreateSequenceDictionary (v4, [43]), samtools’ v1.3 faidx function [44] and STAR’s genomeGenerate option together with an Augustus-derived GFF3 annotation [45] provided with the v.2.0 assembly. The resulting sorted bam alignment files were used for measuring RNA-seq digital gene expression using the provided GFF3 gene annotation and the featureCounts tool in the SubRead package [46].

Homologs in *A. thaliana* were found based on protein homology (TAIR10 v.20101214, https://www.arabidopsis.org/download_files/Proteins/TAIR10_protein_lists/TAIR10_pep_20101214), with the blastp mode of blastall (NCBI v.2.2.26) used to find the closest matching thaliana protein to each Augustus-predicted *I. purpurea* protein sequence with an upper limit p-value for reporting a homolog of 1e-6.

### Differential expression analysis

We used Edgr’s filterByExpr function with the default settings (minimum 10 counts-per-million per gene) to filter the gene set to 25,534 genes [47] We calculated normalization factors that scale each sample by raw library size with Edgr’s calcNormFactors function [48–50]. Read counts were transformed with the voom function in Limma and we used Limma to run linear models that estimated differential expression between samples. [51–53].

We conducted the following comparisons for differential expression:

1. Identifying genes with plastic responses by comparing non-selected control lines in herbicide spray (n=16) to non-selected control lines not sprayed with herbicide (n=15). This comparison identifies plasticity in the original population before selection but note that these lines were grown in lab conditions for three generations, so drift or lab selection could affect our observations of plasticity. We also conducted these comparisons within the two timepoints tested, so we compared leaves collected 8 hours after spray (n=8) to leaves from unsprayed plants collected at the same time (n=8) and leaves collected 32 hours after spray (n=8) to leaves from unsprayed plants collected at the same time (n=7).
2. Identifying genes with evolved responses after selection for herbicide resistance by comparing resistance selection lines not sprayed (n=15) to original lines not sprayed (n=15).
3. Identifying genes with evolved expression responses in the herbicide environment after selection for herbicide resistance by comparing resistance selection lines in spray conditions (n=15) to original lines in spray conditions (n=15).

We conducted GO enrichment with PANTHER on pantherdb.org [54] using the Fisher’s exact test with a false discovery rate analysis. We compared genes that had increased expression in herbicide (3,789 annotated genes) or decreased expression (N annotated genes) to all expressed genes from the sample (12,702 annotated genes).

All code is available at https://github.com/emjosephs/morning-glory and archived at

## Results

### Plastic and evolved expression responses to herbicide

We measured plastic responses to herbicide in control plants that had not been artificially selected for herbicide resistance (Figure 1). Morning glory leaf gene expression responded plastically to herbicide, both 8 and 32 hours after herbicide application. Out of the 27,037 genes tested, 3,519 genes had higher expression and 2,841 genes had lower expression 8 hours after spray (FDR < 0.1) and 5,732 genes had higher expression and 5,285 genes had lower expression 32 hours after spray (FDR < 0.1). The log fold change of expression changes caused by spray was correlated across the 8 and 32 hour measurements (cor coeff = 0.614, p < 0.001, Figure S1). There were only 137 genes with significant (FDR < 0.1) plastic responses to spray at both time points where the direction of response differed between the 8 hour and 32 hour time points. Because of the similarity of response at 8 and 32 hours after spray, we pooled data for future analyses.

**Figure 1:**
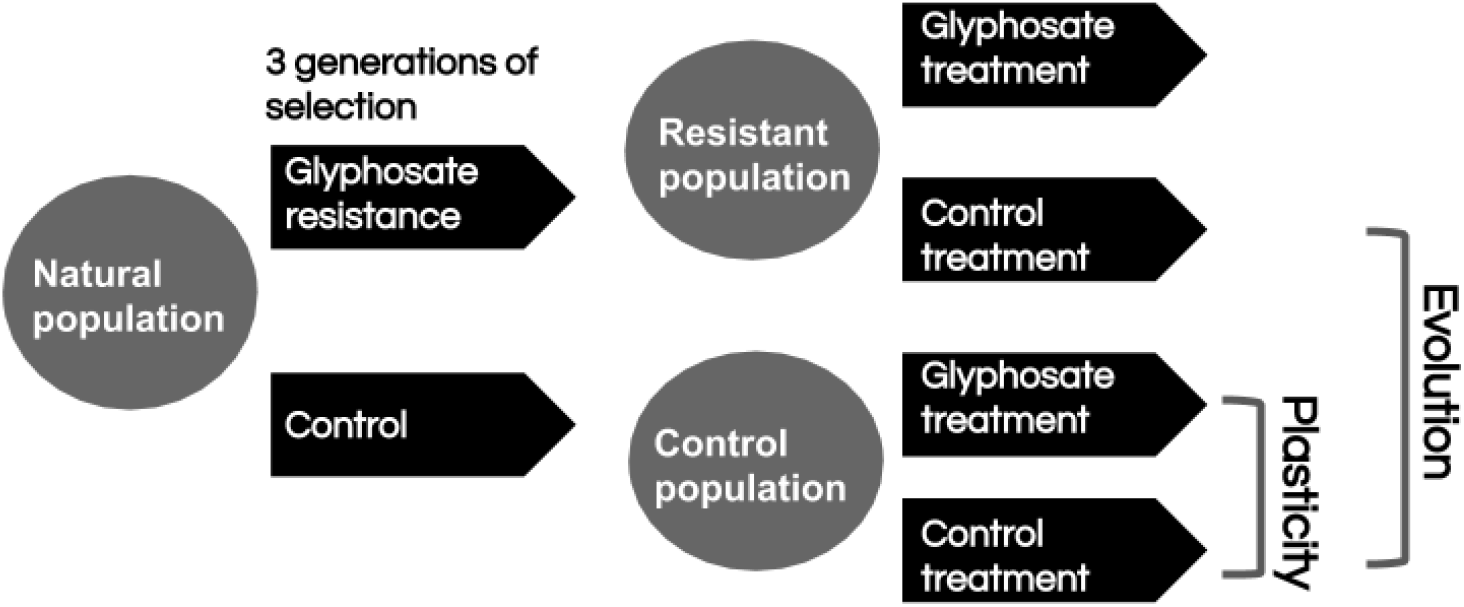
Conceptual figure of the experimental design. This figure shows the breeding scheme that was used to generate the plants used in this study. Plastic expression changes were measured by comparing gene expression in control plants from a control treatment and from the glyphosate treatment. Evolved expression changes were used by comparing control treatment plants from the control and resistance populations.

When data from both time points was pooled to estimate plastic responses to spray in general, 5,734 genes had lower expression in response to pesticide spray and 6,171 had increased expression in response to spray (FDR < 0.1, Figure 2). Genes with increased expression after spray were enriched for a number of GO biological processes at FDR < 0.01, including a number of terms related to abiotic and biotic interactions (Table S1). In contrast, genes with reduced expression after spray were enriched for GO terms related to photosynthesis (Table S2).

**Figure 2:**
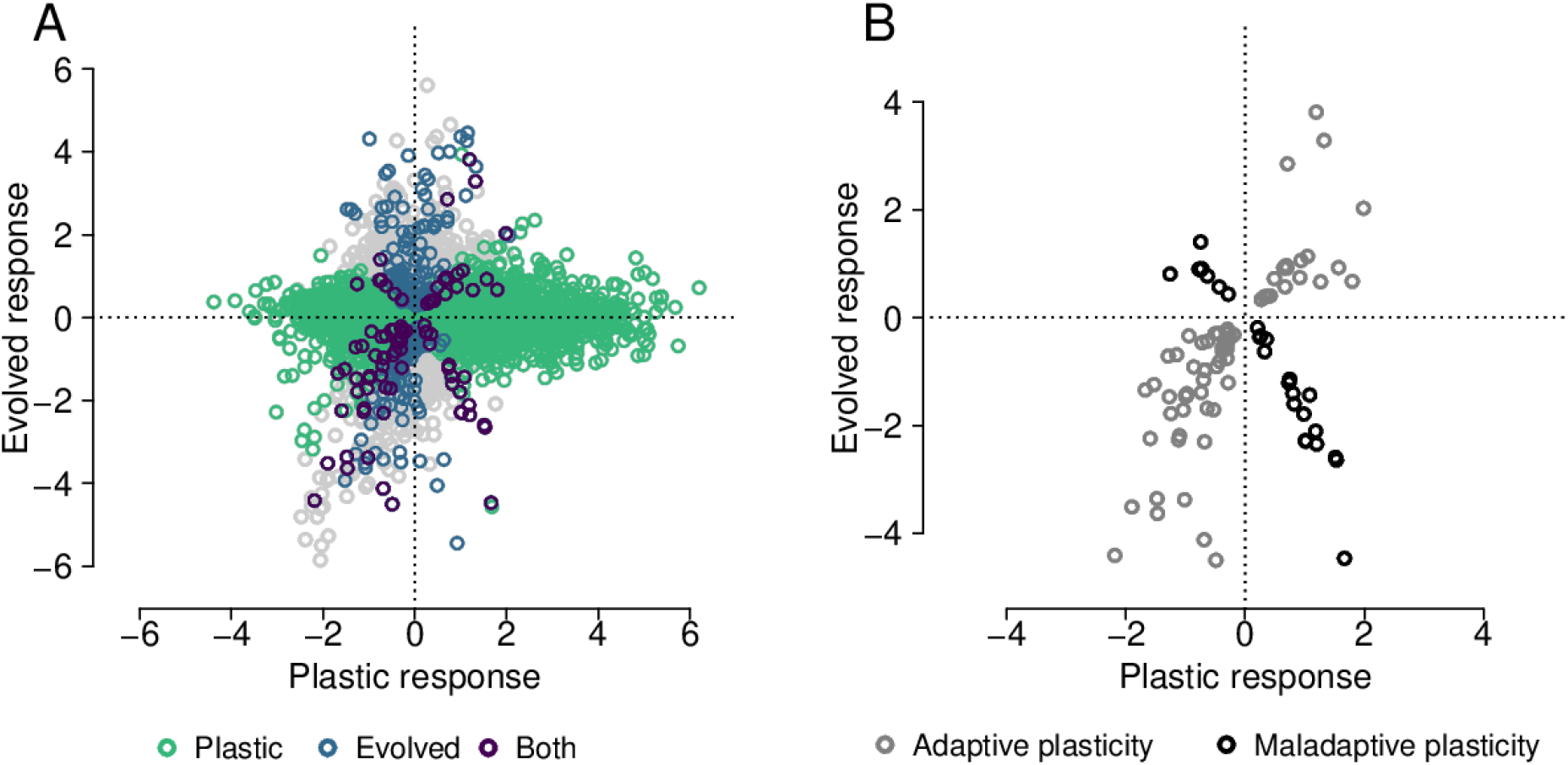
Plastic and evolved changes in expression: A) The x axis shows plastic responses to glyphosate, specifically the log fold change in expression between sprayed and nonsprayed conditions where positive values indicate increased expression in herbicide spray. The y axis shows the log fold difference in expression between lines selected for increased resistance to herbicide and lines that were not selected (‘original lines’), where positive values indicate increased expression in the resistance selection lines. Each point represents one gene, colored by whether they have significant (FDR < 0.1) plastic expression responses, evolved expression responses or both. B) The same as (A), but only genes with significant responses to selection and treatment are shown.

Gene expression in *I. purpurea* leaves responded to selection for resistance (Figure 2). After three generations of selection for herbicide resistance, 166 genes showed decreased expression and 133 genes showed increased expression (FDR < 0.1, Figure 2A). These genes were not significantly enriched for any GO terms.

### The prevalence of adaptive and maladaptive plasticity

There were 94 genes that had both significant plastic responses to spray and evolved responses to selection for resistance in unsprayed conditions (FDR < 0.1). Within these 94 genes, 68 had plastic responses in the same direction as evolved responses, consistent with adaptive plasticity (Binomial p = 1.7 × 10^−5^, Figure 2B). Overall, with the majority of genes showing plastic responses in the same direction as evolved responses, these results show that more of the expression responses in response to herbicide are adaptive than maladaptive. We only looked at selection in non-sprayed conditions to avoid confounding selective response and plasticity [55].

The pattern of adaptive plasticity was also evident for gene expression in samples collected at 8 hours after spray, where 44 out of 56 genes with plastic and evolved responses showed adaptive plasticity (binomial p = 2×10^−5^), and marginally significant at 32 hours after spray, where 44 of 71 genes with significant adaptive and plastic responses exhibited adaptive plasticity (binomial p = 0.057, Figure S2A, B). While there were more genes that showed evidence of either maladaptive or adaptive plasticity 32 hours after spray, the directionality of the response were broadly consistent, with only one of the genes changing direction from adaptive after 8 hours to maladaptive after 32 hours.

### Specific genes showing adaptive and maladaptive plasticity for expression

It was possible to annotate 41 of the 68 genes with evidence of adaptive plasticity based on homology to *Arabidopsis thaliana* (Table S3). Potential genes of interest included the homolog of ATR7, which is associated with oxidative stress tolerance [56] and PRL1, which is associated with growth and immunity [57]. However, many genes identified as having adaptive plasticity did not have homologs with obvious links to processes important for herbicide or other stress responses (Table S3).

We were able to identify homologs from *A. thaliana* for 24 of the 26 genes with evidence of maladaptive plasticity. Many of the genes that show increased plastic expression during spray but decreased expression after selection for resistance were homologs of three *A. thaliana* leucine-rich repeat protein kinase family proteins (Figure 3, Table S3). Leucine-rich repeat protein kinases are often involved in responses to stress [58–61]. Many of these genes were located near each other in the *I. purpurea* reference genome, suggesting that they are recent tandem duplicates. The observed patterns of expression changes suggest that selection for herbicide resistance reduced the expression of these genes, providing a potential future path for understanding the mechanisms of adaptation to herbicides.

**Figure 3:**
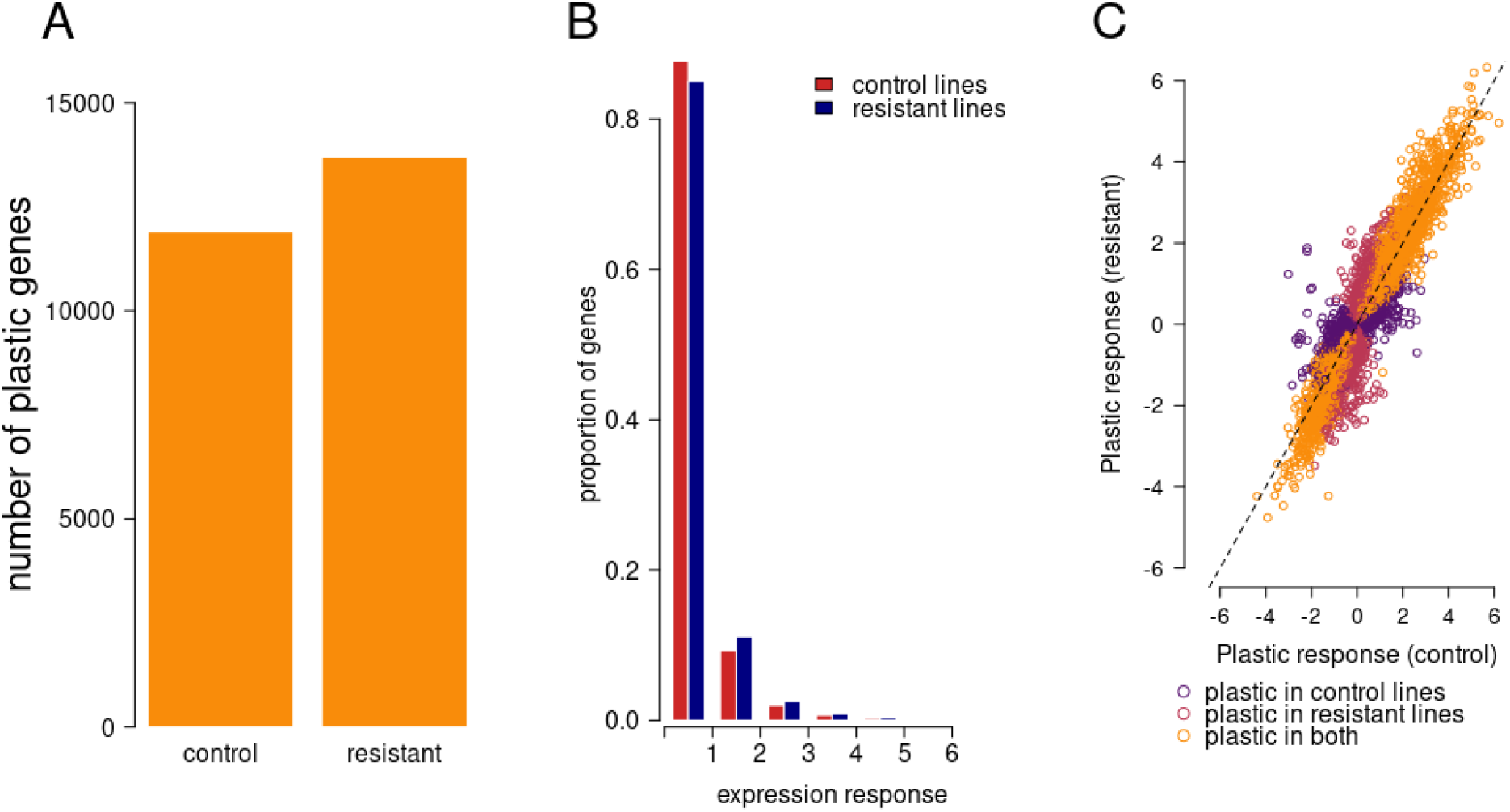
A) The number of genes showing plastic expression differences in response to herbicide at FDR < 0.1 in control and selected lines B) A histogram of gene expression responses to herbicide, measured as the absolute value of the log 2 fold change of expression measured with and without herbicide. C) Plastic expression in control lines and lines selected for resistance. Response is shown on a log2 scale and the x=y line is plotted with a dashed black line. Points are colored by whether they show a significant plastic response to spray in resistant and control lines. Genes without a significant response in either set of lines are not shown.

### Selection for herbicide resistance increases plasticity

If the plastic expression changes that occur in response to spray are adaptive, selection for increased herbicide resistance should increase the extent and magnitude of expression changes in response to herbicide. Consistent with this prediction, more genes had plastic expression responses to herbicide after selection for resistance compared to control lines (FDR < 0.1, Figure 3A). Similarly, 59% of all genes studied had a greater expression response to herbicide after selection for resistance than in control lines (Figure 3B). However, across all genes, plastic responses to selection were correlated in control and resistant lines, suggesting that selection for resistance did not drastically alter patterns of plasticity (Figure 3C, Figure S3).

## Discussion

Here, we have shown that the gene expression plasticity of *Ipomoea purpurea* after glyphosate application shows patterns consistent with being adaptive. The plastic gene expression changes we identified generally align in the same direction as evolved changes after artificial selection for increased herbicide resistance. These results differ from those of other systems, where maladaptive gene expression plasticity appears to be more common [22–25]. There are a number of potential explanations for this difference. First, the population studied here may exist in a set of conditions where plasticity is predicted to be adaptive [1,8,62,63]. For example, the ancestors of the lines used in this study were collected in 2000, after experiencing intermittent glyphosate application for at least 8 years [64], so fluctuating selection from herbicide use may have selected for plastic responses to herbicide. In addition, tolerance and resistance to glyphosate were present in *I. purpurea* before the use of glyphosate, suggesting that these plastic traits may be pleiotropic with plastic responses to other stresses [65]. While in agricultural settings, herbicide is generally applied once per generation, the timing of application could vary year-by-year relative to weed developmental timing. If the physiological effects of herbicidediffer across development, this treatment could contribute to fluctuating selection, a process that can select for adaptive plasticity [5].

There are a number of potential consequences of widespreadadaptive plasticity. Adaptive plasticity could facilitate adaptation to new conditions by allowing populations to persist long enough for adaptation to occur but it could also hinder local adaptation by masking genetic variation from selection [5]. Which process occurs depends on how close the plastic response is to the phenotypic optimum [5,8]. In this study, despite evidence of potential adaptive gene expression plasticity, the population was still able to evolve increased resistance to herbicide [35], suggesting that the plastic responses do not completely move plants to the optimum trait value. Interestingly, the results here suggest that further adaptation to herbicide treatment increased the number of genes with expression plasticity and the magnitude of this plasticity.

It is unclear if and how the results of this studywould change if evolutionary changes in expression were evaluated in natural populations that have evolved resistance instead of in artificially selected populations. The responses to selection that occur in experimental evolution will differ from those in natural populations [66]. An additional limitation for all studies in non-model systems is that we are reliant on GO terms for homologs in *Arabidopsis thaliana*, a distantly related species. A lack of gene or functional conservation will erode our ability to make conclusions about gene function from this data.

There are also limitations to using leaf gene expression as a trait in studies of plasticity. First, leaves are made up of multiple tissues, so variation in different tissue types, instead of direct increases in the transcription of genes within cells, could drive changes in gene expression [67]. Second, the expression level of different genes is not independent and observations of many genes changing in expression could stem from one or a few trans-regulatory factors [68]. Incorporating information about how regulatory networks respond to environmental changes and stresses will be an important next step to understanding the links between selection, plasticity, and adaptation to stressful environments.

This is the first paper to directly investigate the adaptive potential of plastic expression changes in response to herbicide. Previous work has found evidence of gene expression changes in response to herbicide, often in genes not previously known to be important for herbicide response [69–74], but these studies did not directly investigate whether plastic responses appeared to be adaptive. One hint at the importance of maladaptive plasticity comes from the observation that a susceptible population of *Amaranthus tuberculatus* increased rapid expression responses to herbicide compared to a naturally resistant population [75], suggesting that plastic expression changes in response to herbicide could be maladaptive in this species. If our work using an artificially evolved population is any guide for resistance evolution in natural populations, it suggests that selection for increased resistance may reduce expression of some genes, perhaps those with environment-sensing functions (e.g. leucine rich repeat protein kinases), while also leading to increased plasticity in other genes.

In summary, we have shown that *I. purpurea* gene expression responds plastically to herbicide application and evolutionarily to selection for resistance. Crucially, plastic expression responses generally go in the same direction as evolved responses, consistent with the idea that this plasticity is adaptive. We also showed that consistent selection for resistance can increase plasticity, suggesting plasticity is a key component of herbicide resistance in *I. purpurea*. All together, this work demonstrates the importance of adaptive plasticity for the evolution of resistance to environmental stressors.

## Acknowledgements

We thank Jim Leebens-Mack for help with assembly and annotation of the *I. purpurea* reference genome, McKena Wilson and Maya Wilson Brown for research assistance, and Yuheng Huang, Matt Osmond, John Stinchcombe, Sophie Buysse, the Whitehead lab, and two anonymous reviewers for helpful suggestions. This work was funded by Michigan State University, the University of Michigan, and USDA NIFA grants 24892 & 28497 to RSB.

## Supplemental figures

**Figure S1:**
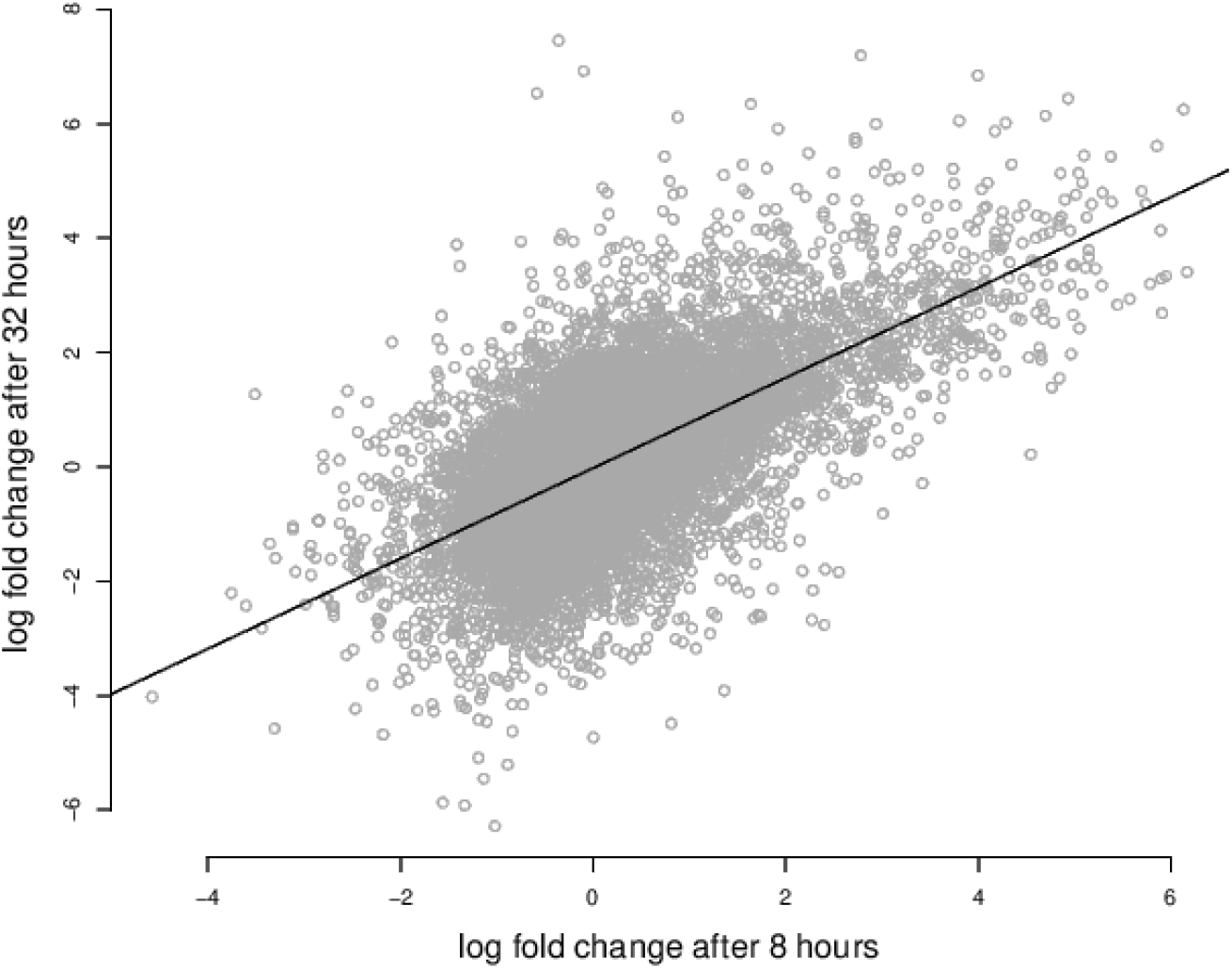
Plastic responses to herbicide spray 8 and 32 hours after treatment. The x axis shows the log-fold difference in expression in sprayed and unsprayed conditions 8 hours after treatment and the y axis shows the same value 32 hours after treatment. The black dotted line shows a line with a slope of 1 and intercept of 0. The correlation coefficient is 0.616. All genes are shown.

**Figure S2:**
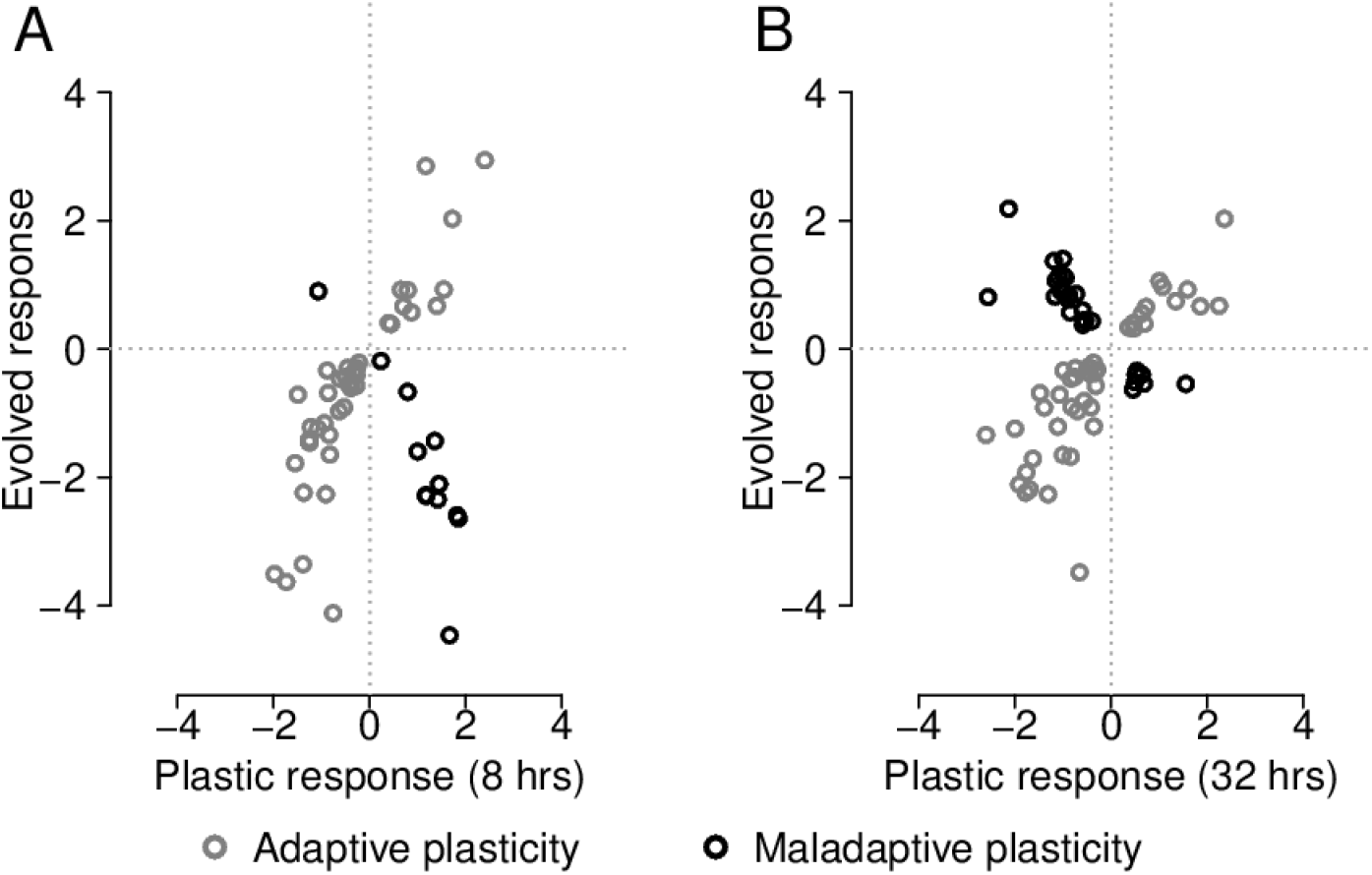
Adaptive and maladaptive expression plasticity at 8 hours (A) and 32 hours (B) after herbicide treatment. In both panels, the x axis shows plastic responses to glyphosate, specifically the log fold change in expression between sprayed and nonsprayed conditions where positive values indicate increased expression in herbicide spray. The y axis shows the log fold difference in expression between lines selected for increased resistance to herbicide and lines that were not selected (‘original lines’), where positive values indicate increased expression in the resistance selection lines. Each point represents a gene and all genes where there was both a significant plastic and an evolved response are shown (FDR < 0.1).

**Figure S3:**
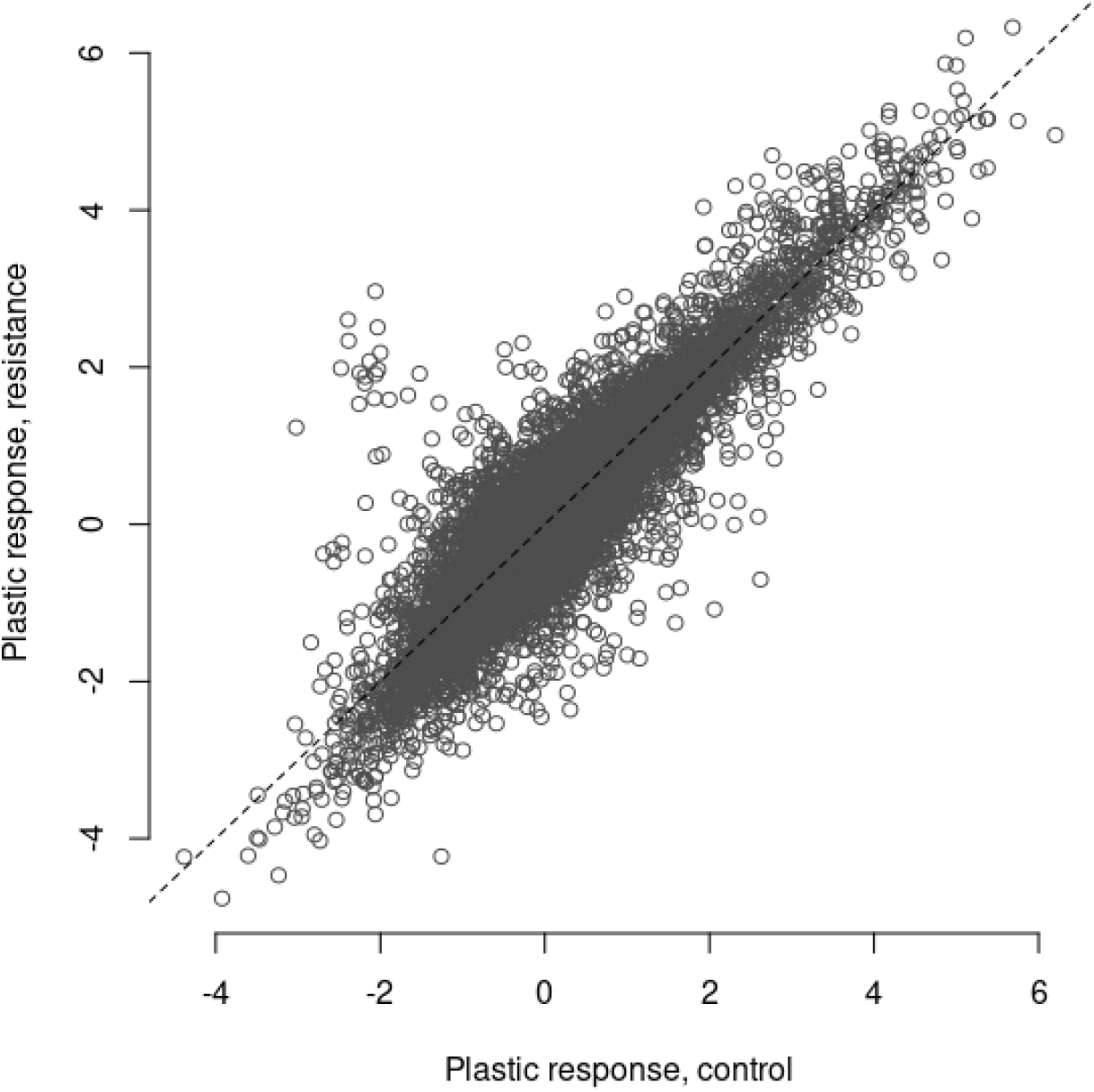
Plastic expression changes in control and resistance lines for all genes. Each point represents one gene, and the y=x line is plotted with a dashed line.

